# Reproducible, flexible and high-throughput data extraction from primary literature: The metaDigitise **R** package

**DOI:** 10.1101/247775

**Authors:** Joel L. Pick, Shinichi Nakagawa, Daniel W.A. Noble

## Abstract

1. Research synthesis, such as comparative and meta-analyses, requires the extraction of effect sizes from primary literature, which are commonly calculated from descriptive statistics. However, the exact values of such statistics are commonly hidden in figures.
2. Extracting descriptive statistics from figures can be a slow process that is not easily reproducible. Additionally, current software lacks an ability to incorporate important meta-data (e.g., sample sizes, treatment / variable names) about experiments and is not integrated with other software to streamline analysis pipelines.
3. Here we present the R package **metaDigitise** which extracts descriptive statistics such as means, standard deviations and correlations from four plot types: 1) mean/error plots (e.g. bar graphs with standard errors), 2) box plots, 3) scatter plots and 4) histograms. **metaDigitise** is user-friendly and easy to learn as it interactively guides the user through the data extraction process. Notably, it enables large-scale extraction by automatically loading image files, letting the user stop processing, edit and add to the resulting data-frame at any point.
4. Digitised data can be easily re-plotted and checked, facilitating reproducible data extraction from plots with little inter-observer bias. We hope that by making the process of figure extraction more flexible and easy to conduct it will improve the transparency and quality of meta-analyses in the future.

## 1 Introduction

In many different contexts, researchers make use of data presented in primary literature. In the fields of ecology and evolution (E&E), these data are most commonly used for comparative and meta-analyses. The use of meta-analysis in E&E in particular, is rapidly growing, not only in terms of the number of meta-analyses (in plant ecology alone the yearly number of published meta-analyses doubled from 2006 to 2012 (20-40) (Koricheva & Gurevitch, 2014)), but also in terms of their size (a recent meta-analysis, for example, included 6440 effect sizes from 175 publications (Noble, Stenhouse & Schwanz, 2018)). Meta-analyses are extremely important in providing a means of quantitatively synthesizing experimental and/or observational studies to evaluate empirical support for fundamental theory in E&E (Gurevitch et al., 2018). These techniques rely heavily on descriptive statistics (e.g. means, standard deviations (SD), sample sizes, correlation coefficients) extracted from primary literature. As well as being presented in the text or tables of research papers, descriptive statistics are frequently presented in figures. For example, 42% of the papers used in a recent meta-analysis presented some or all of the required data in figures (Noble, Stenhouse & Schwanz, 2018). These data need to be manually extracted using digitising programs.

Although there are several tools that extract data from figures, including both standalone programs and R packages (reviewed in Table 1), these tools do not cater to the general needs of meta-analysts for four main reasons (here we focus on meta-analysis, although many points apply to extraction for comparative analysis). First, although meta-analysis is an important tool in consolidating the data from multiple studies, many of the processes involved in data extraction are opaque and difficult to reproduce, making extending or replicating studies problematic. Having a tool that facilitates reproducibility in meta-analyses will increase transparency and aid in resolving the reproducibility crises seen in many fields (Peng, Dominici & Zeger, 2006; Peng, 2011; Parker et al., 2016). Second, digitising programs do not allow the integration of metadata at the time of data extraction, such as experimental group or variable names, and sample sizes. This makes the downstream calculations laborious, as information has to be added later, typically using different software. Third, existing programs do not import sets of images for the user to systematically work through. Instead they require the user to manually import images and export the resulting digitised data into individual files one-by-one. These data often subsequently need to be imported and edited using different software. Finally, digitising programs typically only provide the user with calibrated *x*,*y* coordinates from imported figures, and do not differentiate between common plot types that are used to present data. Consequently, a large amount of additional data manipulation is required, that is different across plots types. For example, in E&E data are commonly presented in plots with means and standard errors or confidence intervals (Figure 1A), from which the user wants a mean and SD for each group presented. From *x,y* coordinates, users must manually discern between mean and error coordinates and assign points to groups. The error then needs to be calculated as the deviation from the mean, and then transformed to SD, according to the type of error presented. Histograms and box plots are also frequently used in E&E to presented data, and whilst their downstream calculations are even more laborious, there are few (if any; see Table 1) tools to extract data from these plot types.

**Table 1:**
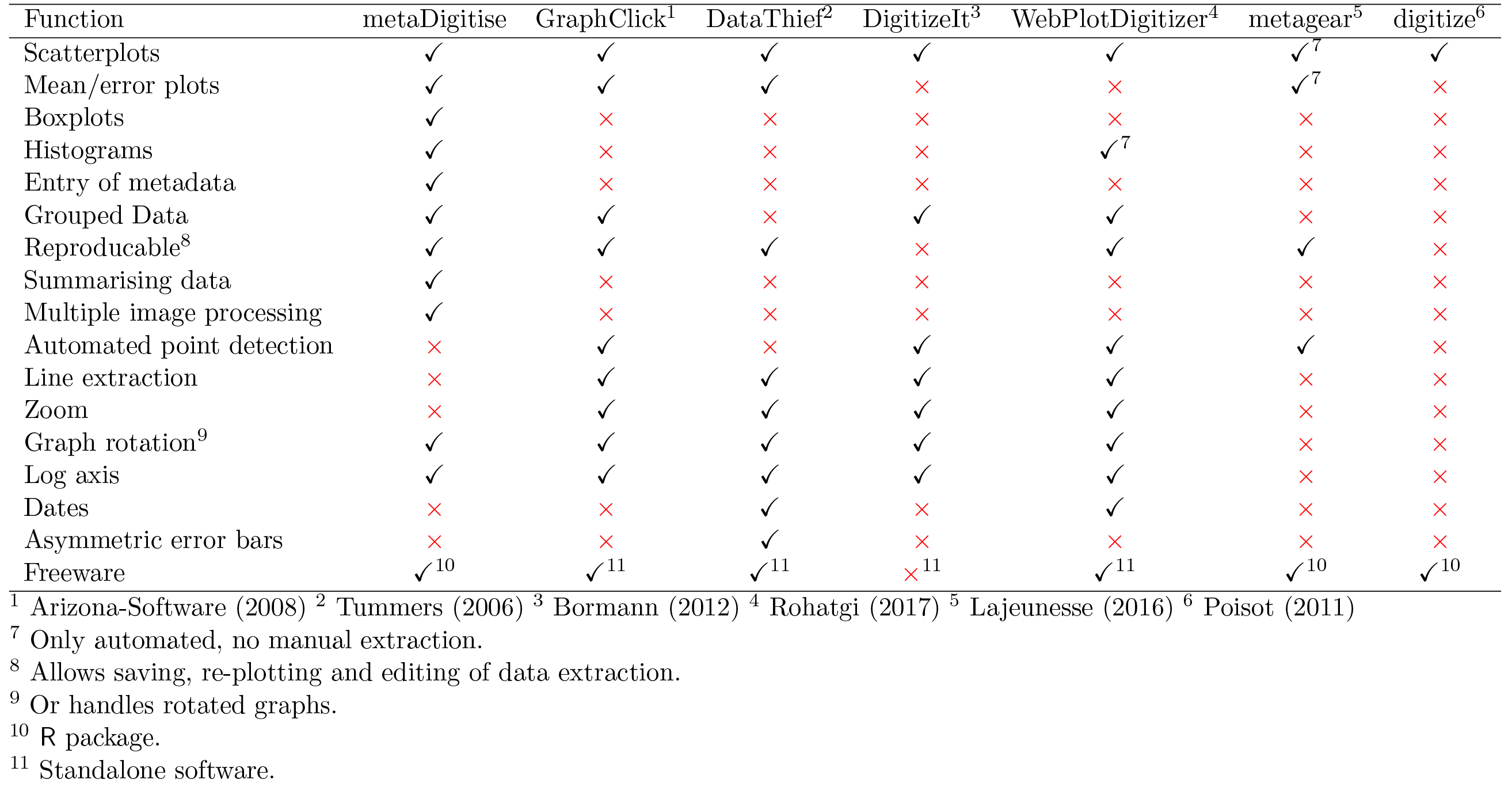
Comparison of functionality between different digitisation softwares.

**Figure 1:**
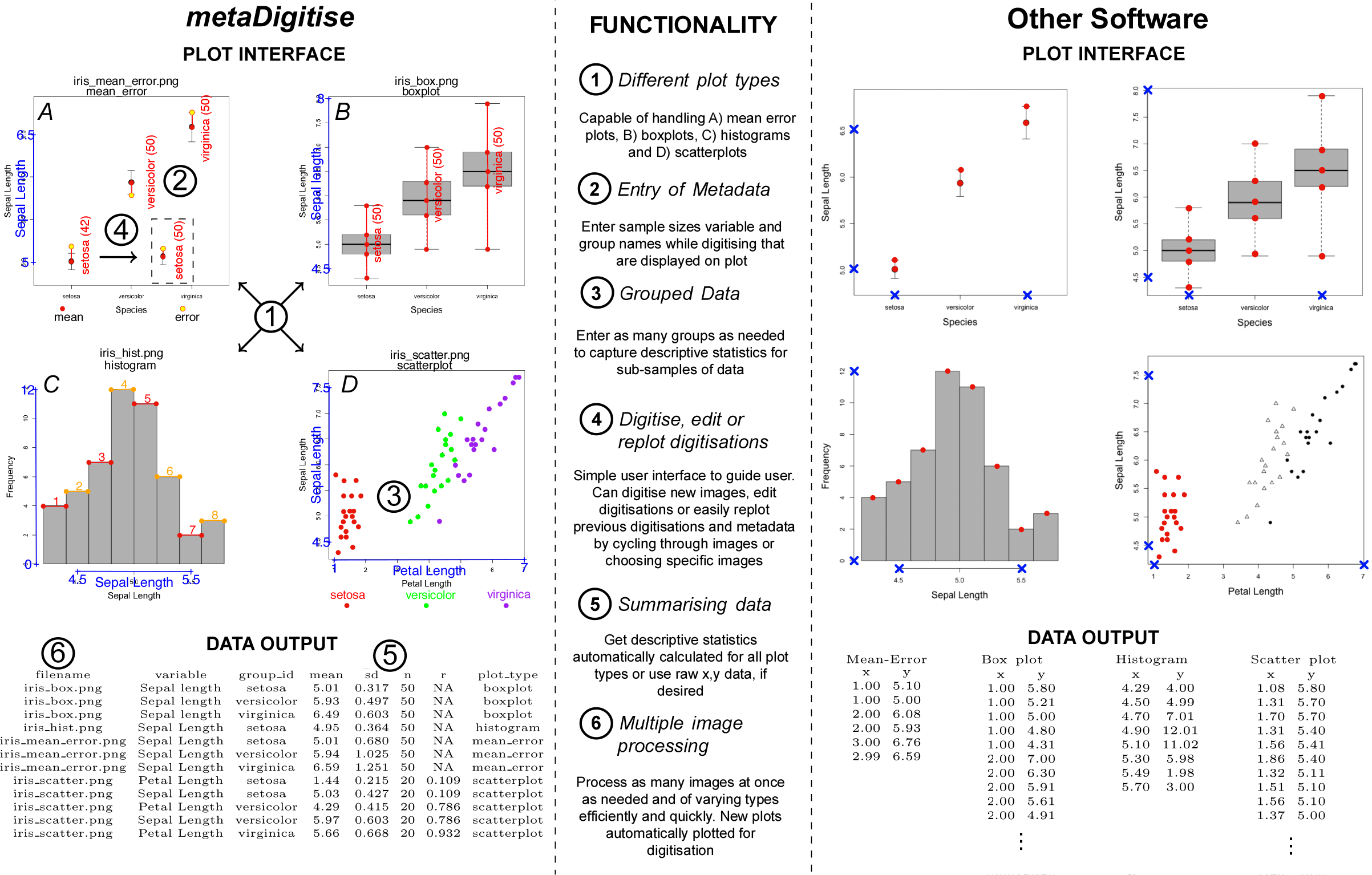
Functionality of **metaDigitise**. Using the iris dataset in R, digitisation of different plot types, A) mean/error plot, B) box plot, C) histogram and D) scatter plot, is shown in **metaDigitise** (left) compared with other common softwares (right). A) and B) are plotted with the whole dataset, C) is just the data for the species *setosa* and D) a subset from all three species. Notable functions of metaDigitise are listed in the center. Other software also perform points 3 and 4 (see Table 1), although these functions are more developed in **metaDigitise**. As shown on the left hand side of the figure, **metaDigitise** clearly displays the stages of the digitisation to aid the transparency of the process, and returns concatenated summary data for all images.

Data extraction from figures is therefore a time-consuming process as existing software does not provide an optimized, reproducible research pipeline to facilitate data extraction and editing. Given the ubiquity of the `R` platform in E&E, and that it hosts the most popular meta-analysis software in E&E (e.g., metafor (Viechtbauer, 2010) and MCMCglmm (Hadfield, 2010)), it is highly likely to be used for some (if not all) stages of the research synthesis process. It is therefore important to have comprehensive, robust and flexible digitisation capabilities in R to make the process of figure extraction more streamline, transparent and easier to reproduce. Here, we present an interactive R package, **metaDigitise** (available on CRAN), which is designed for large scale, reproducible data extraction from figures, specifically catering to the the needs of meta-analysts. To this end, we provide tools to extract data from common plot types in E&E (mean/error plots, box plots, scatter plots and histograms, see Figure 1). **metaDigitise** operates within the `R` environment making data extraction, analysis and export more streamlined. The necessary calculations are carried out on calibrated data immediately after extraction so that comparable descriptive statistics can be obtained quickly. Summary data from multiple figures is returned into a single data frame which can be can easily exported or used in downstream analysis within `R`. Completed digitisations are automatically saved for each figure, meaning users can redraw their digitisations (along with metadata) on figures, make corrections and access calibration and processed (i.e., summarised) data. This makes sharing figure digitisation and reproducing the work of others simple and easy, and allows meta-analyses to be updated more efficiently.

## 2 metaDigitise and Reproducibility

The **metaDigitise** package has one main function, `metaDigitise()`, which interactively takes the user through the process of extracting data from figures (see Supplementary Material S1 for a full tutorial). Running `metaDigitise()` presents the user with three options; ‘Process new images’, ‘Import existing data’ or ‘Edit existing data’, which can be used during and after digitisation to execute a range of functions (see Figure 1 - ‘Processing images’ is discussed in Section 3, and ‘Editing’ and ‘Importing’ in Section 4). `metaDigitise()` works on a directory containing images of figures copied from primary literature, in .png, .jpg, .tiff, .pdf format, specified to `metaDigitise()` through the `dir` argument. `metaDigitise()` recognizes all the images in the given directory and automatically imports them one-by-one, allowing the user to extract the relevant information about a figure as they go. Figures can be organised in different ways for a project, but we would recommend having all figures for one project in a single directory with an informative and unambiguous naming scheme (e.g. paper_figure_trait.png). This expedites digitisation by preventing users from having to constantly change directories and / or open new images.

The data from each completed image is automatically saved as a `metaDigitise` object in a separate .RDS file to a `caldat` folder that is created within the parent directory when first executing `metaDigitise()`. These files enable re-plotting and editing of images at a later point (see below). When run, `metaDigitise()` also identifies the images within a directory that have been previously digitised and only imports new images to process. The data of all images is then automatically integrated into the final output. This means that all figures do not need to be extracted at one time and new figures can be added to the directory as the project develops.

The complete digitisation process can be reproduced at a later stage, shared with collaborators and presented as supplementary materials for a publication, regardless of the computer it is run on. To update an analysis, new figures can simply be added to the directory and `metaDigitise()` run to incorporate the new data.

## 3 Image Processing

Selecting ‘Process New Images’, after running `metaDigitise()`, starts the digitisation process on images within the directory that have not previously been digitised. For all plot types, `metaDigitise()` requires the user to calibrate the axes in the figure, by clicking on two known points on the axis in question, and entering the value of those points (Figure 1). `metaDigitise()` then calculates the value of any clicked points in terms of the figure axes. This is based on the calibration used in the **digitize** R package (Poisot, 2011). For mean/error and box plots, only the y-axis is calibrated (Figure 1), assuming the x-axis is redundant. For scatter plots and histograms both axes are calibrated (Figure 1).

Calibration of points in figures from older, scanned publications can be problematic, as the figures may not be perfectly orientated. `metaDigitise()` allows users to rotate the image (Figure S2A,B). Furthermore, mean/error plots, box plots and histograms, may be presented with horizontal bars. `metaDigitise()` assumes that bars are vertical, but allows the user to flip the image to make the bars are vertical (Figure S2C,D). **metaDigitise** also allows back calculation of data presented on log axes.

**metaDigitise** recognises four main types of plot; Mean/error plots, box plots, scatter plots and histograms (Figure 1). All plot types can be extracted in a single call of `metaDigitise()` and integrated into one output. Alternatively, users can process different plot types separately, using separate directories. All four plot types are extracted slightly differently (outlined below). Upon completing all images, or quitting, either summarised or calibrated data is returned (specified by the user through the `summary` argument). Summarised data consists of a mean, SD and sample size, for each identified group within the plot (should multiple groups exist). In the case of scatter plots, the correlation coefficient between x and y variables within each identified group is also returned. Calibrated data consists of a list with slots for each of the four figure types, containing the calibrated points that the user has clicked. This may be particularly useful in the case of scatter plots.

### 3.1 Mean/Error and Box Plots

`metaDigitise()` handles mean/error and box plots in a very similar way. For each mean/box, the user enters group name(s) and sample size(s). If the user does not enter a sample size at the time of data extraction (if, for example, the information is not readily available) a SD is not calculated. Sample sizes can, however, be entered at a later time (see next section). For mean/error plots, the user clicks on an error bar followed by the mean. Error bars above or below the mean can be clicked, as sometimes one is clearer than the other. `metaDigitise()` assumes that the error bars are symmetrical. Points are displayed where the user has clicked, with the error in a different colour to the mean (Figure 1A). The user also enters the type of error used in the figure: SD, standard error (SE) or 95% confidence intervals (CI95). For box plots, the user clicks on the maximum, upper quartile, median, lower quartile and minimum. For both plot types, the user can add, edit or remove groups while digitising for when finished. Three functions, `error_to_sd(), rqm_to_mean() and rqm_to_sd(),` that convert different error types to SD, box plot data to mean and box plot data SD, respectively, are also available in the package (see supplements for further details of these conversions).

### 3.2 Scatter plots

Users can extract points from multiple groups from scatter plots. Different groups are plotted in different colours and shapes to enable them to be distinguished, with a legend at the bottom of the figure (Figure 1D). Mean, SD and sample size are calculated from the clicked points, for each group. Data points may overlap with each other making it impossible to know whether points have been missed. This may result in the sample size of digitised groups conflicting with what is reported in the paper. However, users also have the option to input known sample sizes directly, if required. Nonetheless, it is important to recognise the impact that overlapping points can have on descriptive statistics, and in particular on sampling variance.

### 3.3 Histograms

The user clicks on the top corners of each bar, which are drawn in alternating colours (Figure 1C). Bars are numbered to allow the the user to edit them. As with scatter plots, if the sample size from the extracted data does not match a known sample size, the user can enter an alternate sample size. The formulas for calculation of mean, SD and sample size are provided in the supplement.

## 4 Importing and Editing Previously Digitised data

**metaDigitise** is also able to re-import, edit and re-plot previously digitised figures. When running `metaDigitise()`, the user can choose to ‘Import existing data’, which returns previously digitised data, from a single figure or all figures. Alternately, the `getExtracted()` function returns the data from previous digitisations, but without user interaction, allowing easier integration into larger scripts. ‘Edit existing data’ allows the user to re-plot or edit information for digitisations that have previously be done. Re-plotting digitisations with all metadata is an important reproducibility feature, as it allows users to see exactly what information has been extracted, as well as making it easy to spot and data extraction errors.

### 4.1 4.1 Adding Sample Sizes to Previous Digitisations

In many cases sample sizes may not be readily available when digitising figures. This information does not need to be added at the time of digitisation. To expedite finding and adding these sample sizes at a later point, `metaDigitise()` has a specific edit option that allows users to enter previously omitted sample sizes. This first identifies missing sample sizes in the digitised output, re-plots the relevant figures and prompts the user to enter the sample sizes for the relevant groups in the figure.

## 5 Software Validation

To evaluate the consistency of digitisation with **metaDigitise** between users, fourteen people digitized sets of 14 identical images created from a simulated dataset (see supplements). We found no evidence for any inter-observer variability in digitisations for the mean (ICC = 0, 95% CI = 0 to 0.029, *p* > 0.999), SD (ICC = 0, 95% CI = 0 to 0.033, *p* > 0.999) or correlation coefficient (ICC = 0.053, 95% CI = 0 to 0.296, *p* = 0.377). There was little bias between digitised and true values, on average 1.63% (mean = 0.02%, SD = 4.9%, *r* = -0.03%) and there were small absolute differences between digitised and true values, on average 2.18% (mean = 0.40%, SD = 5.81%, r = 0.33%) across all three descriptive statistics. SD estimates from digitisations are clearly most error prone. The mean absolute differences for each plot type clearly show that this effect is driven by extraction from box plots and histograms (% difference; box plot: 15.81, histogram: 5.21, mean/error: 1.50, scatter plot: 0.43). SD estimation from box plot descriptive statistics is known to be more error prone, especially at small sample sizes (Wan et al., 2014).

We also used simulated data to test the accuracy of digitisations with respect to known values (see supplements). **metaDigitise** was very accurate at matching clicked points to their true values essentially being perfectly correlated with the true simulated data for both the *x*-variable (Pearson’s correlation; *r* > 0.999, *t* = 2137.4, *df* = 78, *p* < 0.001) and *y*-variable (*r* > 0.999, *t* = 1897.8, *df* = 78, *p* < 0.001) in scatterplots.

## 6 Limitations

Although **metaDigitise** is very flexible and provides functionality not seen in any other package, there are some functions that it does not perform (see Table 1). Notably **metaDigitise** lacks automated point detection. However, from our experience, manual digitising is more reliable and often equally as fast. Given the variation in image quality, calibration for automatic point detection needs to be done for each figure individually. Additionally, auto-detection often misses points which then need to be manually added. Based on tests of **metaDigitise** (see above), figures can be extracted in around 1-2 minutes, including the entry of metadata. As a result, we do not believe that current automated point detection techniques provide substantial benefits in terms of time or accuracy. Indeed, in a recent project developing automated point extraction techniques, only 15/136 (11%) of studies screened contained figures suitable for the presented method, and in only 12/27 (44%) of the resulting figures was the data correctly extracted (Hartgerink & Murray-Rust, 2017).

**metaDigitise** also (currently) lacks the ability to zoom in on figures. Zooming may enable users to gain greater accuracy when clicking on points. However, from our own experience (see results above), with a reasonably sized screen accuracy is already high, and so relatively little gain is to be had from zooming in on points.

In contrast to some other packages **metaDigitise** does not extract lines from figures. Although line extraction is not generally necessary in comparative and meta-analysis, outside of these fields researchers may need to extract parameters of a line from a figure. Should a user like to extract lines with **metaDigitise**, we would recommend extracting data as a scatter plot, and clicking along the line in question. A model can then be fitted to these points (accessed by choosing to return calibrated rather than summary data) to estimate the parameters needed.

## 7 Conclusions

Increasing the reproducibility of figure extraction for meta-analysis and making this laborious process more streamlined, flexible and integrated with existing statistical software will go a long way in facilitating the production of high quality meta-analytic studies that can be updated in the future. We believe that **metaDigitise** will improve this research synthesis pipeline, and will hopefully become an integral package that can be added to the meta-analysts toolkit.

## Acknowledgments

We thank the I-DEEL group and colleagues at UNSW for for testing, providing feedback and digitising including: Rose O’Dea, Fonti Kar, Malgorzata Lagisz, Julia Riley, Diego Barneche, Erin Macartney, Ivan Beltran, Gihan Samarasinghe, Dax Kellie, Jonathan Noble, Yian Noble, Elena Noble and Alison Pick. J.L.P. was supported by a Swiss National Science Foundation Early Mobility grant (P2ZHP3_164962), D.W.A.N. was supported by an Australian Research Council Discovery Early Career Research Award (DE150101774) and UNSW Vice Chancellors Fellowship and S.N. an Australian Research Council Future Fellowship (FT130100268).

## Author Contributions

J.L.P. and D.W.A.N. conceived the study and J.L.P., S.N. and D.W.A.N. developed the idea. J.L.P. and D.W.A.N. developed the R-package. J.L.P. and D.W.A.N. wrote the first draft of the paper and J.L.P., S.N. and D.W.A.N. contributed substantially to subsequent revisions of the manuscript and gave final approval for publication.

